# A Systematic Approach Toward Implementing Machine Learning Techniques to Analyze Gut Microbiome Data

**DOI:** 10.64898/2026.04.22.720178

**Authors:** Anvi Taada, Ava George, Dhatrisri Biruduraju, Emily Lu, Isha Singh, Khushi Chhajer, Madeline Wang, Tanvi Pentela, Sahar Jahanikia

**Author notes:** Corresponding author: Sahar Jahanikia. All authors contributed equally to this work. Author order was determined by names in alphabetical order.

## Abstract

This study investigates the relationship between the gut microbiota and specific diseases. Data was collected from the Human Gut Microbiome Atlas, which examines regional variations across 20 countries on five continents, categorizing microbial species by taxonomy, from genus to species. The Atlas provides color-coded phylum classifications, numerical species counts within the same genus, and an analysis of dysbiosis-related associations with 23 diseases, as well as region-enriched species. The data stratified samples into distinct categories such as westernized, non-westernized, cancerous, and non-cancerous. The findings demonstrate that tree-based ensemble methods, such as Bagging and Boosting prediction methods, achieved the highest accuracies across all categories due to their robustness in handling the complex, high-dimensional data. The XGBoost model yielded the strongest predictive performance, achieving 91% accuracy for westernized cancer-associated samples, 84% accuracy for non-westernized cancer-associated samples, 92% accuracy for westernized samples, and 78% for non-westernized samples. Additionally, advanced topological data analysis was used to assess the global structure and underlying patterns within the dataset.

**Importance:** This research aims to connect gut microbiome composition to diseases using global datasets from the Human Gut Microbiome Atlas. The goal was to evaluate how accurately different machine learning algorithms could classify microbiota species and diseases and predict disease associations by comparing westernized and non-westernized populations, including both cancerous and noncancerous groups. These findings can contribute to the future creation of population-specific and disease-specific microbial models.

## 1. INTRODUCTION

Different disorders are influenced in the gut-brain axis, which is the communication system between the gastrointestinal tract and nervous system, by gut microbiota. Gut microbiota supports neuroinflammation, blood-brain barrier permeability, and cognitive function (1). Existing evidence shows the microbiota’s influence across Alzheimer’s, Parkinson’s, stroke, mental health disorders, autism, obesity, diabetes, and inflammatory disease (2). Recent studies have identified experimental models to assess microglial activity, neuroinflammatory signatures, and cognitive outcome (3). Prior work also excels at identifying disease-specific microbial signatures and linking dysbiosis to key pathological mechanisms (4). In-vitro models have been developed to measure the cognitive effect of gut bacteria through changes in gene expression, exhibiting the gut microbiome’s direct influence on the gut-brain axis (5).

Global gut microbiome datasets provide information regarding the intersection of human gut microbial composition, taxonomic rank of organisms found, geographical location, and disease. The data consists of gut and oral samples from both diseased and healthy cohorts to determine positive or negative associations of a certain bacteria to a disease (6). In addition, region enrichment demonstrates how environmental factors affect the gut microbiome, resulting in different diseases having different prevalence in varying regions (7). This allows for categorization of diseases that share similar trends in diseases and microbial species that are present, which fall into Westernized and Non-Westernized categories, highlighting how diet and regional habits shape gut dysbiosis. Machine learning and artificial intelligence have become powerful tools in analyzing the gut microbiome through establishing patterns and correlations (8). The applications of machine learning in this study are centered around identifying the type of disease and its presence when the algorithm is provided with the microbial species present, the geographical origin of the sample, and a disease association score. Machine learning techniques for disease detection from the gut microbiome include supervised learning methods like random forest and neural networks to discover patterns with the data (8). However, the large feature space can often lead to overfitting and temporal changes in the disease states. Bagging and boosting, which both utilize decision tree classification, performed the best because they were best suited to the size and complexity of the datasets. TSNE and U-MAP were used for dimensionality reduction in converting the data from four dimensions to two in order to prepare it for Topological Data Analysis (TDA). Prior work does not rely on diverse multi-omics data and does not focus on cross-disease microbial mechanisms, limiting applications in clinical settings. Techniques such as advanced TDA that can capture higher-order structure and nonlinearity are underutilized (9). TDA allows the study of the geometric shape or structure of a dataset for visual representation of its patterns and clusters (10). Kepler Mapper, an advanced TDA tool, is utilized to generate interactive visualizations of the data based on microbial species, genus, country, disease, and association score. The mapper graphs, in addition to uncovering hidden subgroup relationships, reflect previously seen disease and country groupings. This study addresses gaps in current work by integrating datasets that examine regional variations of microbial species by disease. It analyzes the gut microbiome project dataset using advanced data analysis techniques to improve the accuracy of the prediction algorithms for identifying gut-related diseases and microbiome associations. This work integrates multi-omics data for functional insights, such as which microbes exert the most influence on disease outcomes and links microbial gene families, pathways, and disease phenotypes using machine learning techniques. Ultimately, it seeks to develop predictive frameworks that use gut microbiome composition to improve early detection of disease.

## 2. BACKGROUND

The gut microbiome acts as a metabolic organ that influences nutrient absorption, immunity, and mental health (11). It communicates with other organs through the gut-brain and gut-liver axes (12). Microbial balance is essential for health (13) as dysbiosis is linked to conditions (14), such as IBD, obesity, cancer, and mental disorders (15), and early microbial shifts may signal disease onset (16). The gut-brain axis links the gut and CNS through neural, immune, metabolic, and hormonal pathways (17), and dysbiosis is associated with neurological and neurodegenerative disorders (18). Microbial metabolites and immune signaling affect neuroinflammation and blood-brain barrier integrity (19) and microglial regulation (20). Short-chain fatty acids influence neuroplasticity, cognition, and memory (21). Because the microbiome is modifiable, it’s a promising target for early detection and treatment of CNS-related diseases (22), and tracking bacterial shifts may support early risk identification (1).

High-dimensional microbiome data require Machine Learning (ML) to detect patterns despite overfitting risk and heavy preprocessing (23). Increasing dimensionality creates sparsity and computational complexity, making models like Random Forest, XGBoost, Neural Networks, and Support Vector Machine (SVM) essential for classification and prediction (24). Support vector machines are supervised learning methods that construct optimal hyperplanes in high-dimensional feature spaces to maximize classification margins (25,26). ML is applied to classify microbial profiles and predict disease outcomes (27). Geographic variation shapes microbiome composition, highlighting the importance of diverse datasets like the Human Gut Microbiome Atlas. Westernized populations show lower diversity than non-Westernized groups (28). Regional disease patterns reflect interactions among environment, genetics, and public health factors (29). Global datasets improve generalizability and reduce bias in microbiome-disease models.

In Alzheimer’s and Parkinson’s, families like Enterobacteriaceae and genera like Escherichia-Shigella are often present. (30). Beneficial taxa like Lachnospiraceae and Ruminococcaceae are associated with depression, anxiety, and autism spectrum disorder (31). Bifidobacteriaceae and Faecalibacterium consistently appear in datasets on depressive disorder and Rett syndrome, suggesting protective roles (32). Genera like Sutterella, Bilophila, and Prevotella are linked to neurodevelopmental and inflammatory disorders such as ADHD and pervasive developmental disorder (33).

Because microbiome data are high-dimensional, dimensionality reduction is essential for visualization and interpretation while preserving structure (34). Uniform Manifold Approximation and Projection (UMAP) preserves local neighborhoods and global relationships in high-dimensional data (34), performing well on complex biological datasets and revealing patterns missed by linear methods like PCA (35). Similarly, T-distributed Stochastic Neighbor Embedding (t-SNE) preserves local similarity by modeling pairwise relationships (36) and is widely used to visualize subgroup patterns within heterogeneous omics data (37). Clustering often follows dimensionality reduction to group similar samples. DBSCAN identifies clusters as high-density regions separated by low-density areas without requiring a preset number of clusters (38). It also accounts for noise and outliers, making it suitable for sparse, heterogeneous biological data. (39). In high-dimensional analyses, DBSCAN detects intrinsic structure without assuming cluster shape or size (40). When combined with nonlinear methods like UMAP or t-SNE, density-based clustering enables exploration of complex biological variation and interpretation of latent microbiome structure. (35).

## 3. METHODOLOGY

The data used in this study came from several available microbiome databases. The main dataset was taken from the Human Gut Microbiome Atlas, which includes gut microbiota data from 20 countries across five continents (Human Gut Microbiome Atlas, n.d.) as shown in **Table 1**. This dataset provides information on microbial classifications at different taxonomic levels, along with details on disease-associated species, region-specific microbes, and patterns related to gut dysbiosis.

**Table 1:** Table Summary of data collected from Human Gut Microbiome Atlas.

| *Country* | *No. of Species* | *No. of Genera* | *No. of Diseases* |
| --- | --- | --- | --- |
| Austria | 91 | 47 | 19 |
| China | 13 | 9 | 11 |
| Denmark | 125 | 56 | 17 |
| Fiji | 134 | 70 | 16 |
| France | 128 | 67 | 18 |
| Germany | 110 | 59 | 17 |
| India | 27 | 19 | 12 |
| Italy | 80 | 48 | 18 |
| Japan | 55 | 28 | 17 |
| Luxembourg | 98 | 49 | 18 |
| Madagascar | 164 | 72 | 18 |
| Mongolia | 169 | 69 | 18 |
| Peru | 234 | 83 | 16 |
| Spain | 120 | 65 | 19 |
| Sweden | 176 | 79 | 18 |
| Thailand | 120 | 57 | 17 |
| United Kingdom | 76 | 45 | 18 |
| United Republic of Tanzania | 204 | 79 | 16 |
| United States of America | 6 | 4 | 8 |
| Total | 582 | 161 | 21 |

Both Table 1 and 2 were collected from the Human Gut Microbiome Atlas. They give a context to which countries and genera were included in the dataset. The tables convey the relationship between genera, countries, specific diseases, and species. Table 2 highlights the number of species and countries that are affected by a specific genus for the top 10 genera.

**Table 2:** Table of the top 10 genus data with the largest number of species collected from Human Gut Microbiome Atlas.

| *Genus* | *No. of Species* | *No. of Countries* |
| --- | --- | --- |
| Unclassified Lachnospiraceae | 44 | 15 |
| Unclassified RF39 Mollicutes | 22 | 6 |
| Bacteroides | 21 | 18 |
| Blautia | 19 | 17 |
| Lachnoclostridium | 18 | 16 |
| Streptococcus | 17 | 15 |
| Unclassified Clostridiales 2 | 16 | 12 |
| Unclassified Ruminococcaceae | 15 | 16 |
| Unclassified Clostridiales 5 | 14 | 15 |
| Roseburia | 12 | 13 |

### 3.1 DATA ANALYSIS AND VISUALIZATION

Exploratory data analysis was performed to examine how categorical predictors explain variation in Disease Association scores. The enrichment values for gut microbiomes on a range of taxonomic levels, including genus through superkingdom, were obtained from the Human Gut Microbiome Atlas. An ANOVA model was used to evaluate the effects of Disease, Genus, Species Name, and Country on Disease Association scores. The model separated the total variance into components explained by each categorical variable. The output provided degrees of freedom (df), sums of squares (Sum Sq), mean square (Mean Sq), F values, and p-values. Degrees of freedom reflect the number of levels within each factor minus one. Sum of squares represents the total variance explained by each predictor while mean square depicts variance per degree of freedom. The F value measures the strength of the effect, with higher values indicating stronger effects and the p-value determines whether the effect is statistically significant.

### 3.2 MACHINE LEARNING

Various machine learning methods, consisting of K-Nearest Neighbors (KNN), Random Forest, clustering using DBSCAN, Gradient Boosting (Boosting), and Neural Networks (NN), were used to identify links among microbiome variables and disease types. The performances of models were evaluated for the classification accuracy of disease by generating heatmaps, histograms, bar charts, and confusion matrices to compare enrichment patterns across countries and taxonomic ranks.

### 3.3 DIMENSIONALITY REDUCTION

Since the dataset consists of (RowxCol), the Dimensionality reduction was performed using Principal Component Analysis (PCA), Uniform Manifold Approximation and Projection (UMAP), and t-distributed Stochastic Neighbor Embedding (t-SNE) to generate two-dimensional representations of the high-dimensional microbiome dataset. PCA was implemented as an initial linear baseline to evaluate global variance structure. UMAP and t-SNE were subsequently applied to standardized feature matrices to better capture nonlinear relationships among microbial taxa, disease categories, and geographic classifications. All resulting embeddings were visualized as two-dimensional scatter plots, with samples color-coded according to disease status, cancer association, and westernized versus non-westernized groupings. Cluster separation, dispersion, and structural coherence were systematically compared across PCA, UMAP, and t-SNE to assess which method most effectively preserved biologically meaningful patterns.

### 3.4 TOPOLOGICAL DATA ANALYSIS

Topological data analysis (TDA) was performed using KeplerMapper with a t-SNE lens, which produced better clusters than UMAP. Multiple Mapper graphs were constructed using varying values of n_cubes and overlap parameters to assess the stability and persistence of topological features across resolutions. The number of cubes (n_cubes) was systematically varied (5, 10, 15, 20, and 25), while the epsilon value was fixed at 0.3, after testing variations in DBSCAN parameters. Changes in topological structure were analyzed as a function of resolution and hyperparameter selection. Distance metrics were also evaluated, including Euclidean and cosine distances, to examine their effects on graph structure.

## 4. RESULTS

### 4.1 EXPLORATORY DATA ANALYSIS

The results in Table 3 display the figures after conducting ANOVA on a linear regression model on the original data. This demonstrates how much each categorical predictor explains variation in the Disease Association, how strong each effect is, and whether each effect is statistically significant. The sum of squares(sum sq) is the highest for the variable species at 6.5675, followed by the 2nd highest, which is the variable Genus at 3.7126. A larger sum sq for Genus and Species means that the different organisms have vastly different disease associations. The Pr values or the P-values confirm this because genus and species are statistically significant, being less than 0.05. A P-value that is less than 0.05 means that the pattern seen was not a random occurrence. The P-value also confirms patterns with disease and disease association, but not country, because country has a p-value greater than 0.05. Thus, Country proves to have no statistical significance with an F-value of approximately 0.43 and a p-value of 0.983, indicating that Country does not help determine disease association, mainly due to almost no variation in Disease Association across different countries. To end with, biological factors of Genus, Species, and even Disease can predict disease association the best.

**Table 3:** ANOVA on Linear Regression 5×5.

|  | *Df* | *Sum Sq* | *Mean Sq* | *F value* | *Pr(>F)* |
| --- | --- | --- | --- | --- | --- |
| *Disease* | 16 | 2.9088 | 0.1818 | 40.6964 | 6.1042e<sup>-120</sup> |
| *Genus* | 124 | 3.7126 | 0.0299 | 6.7023 | 4.0638e<sup>-95</sup> |
| *Species* | 274 | 6.5675 | 0.02397 | 5.3655 | 2.2480e<sup>-136</sup> |
| *Country* | 18 | 0.0343 | 0.0019 | 0.4270 | 9.8287e<sup>-1</sup> |
| *Residuals* | 4885 | 21.8226 | 0.0045 | NA | NA |

The hierarchical clustering heatmap shows the correlation between gut microbiome species and different diseases. The different colors represent the relative abundance, or association strength, for the microbial species and each of the diseases. The darker red shades indicate a stronger positive correlation, whereas blue hues indicate a negative association, and white areas indicate little to no association.

Table 4 highlights the strongest positive associations found between microbial species and certain diseases, indicated by the darkest red coloring in **Figure 1**. This exploratory data analysis allows for the pinpointing of specific biomarkers to focus on for certain diseases and treatments.

**Figure 1:**
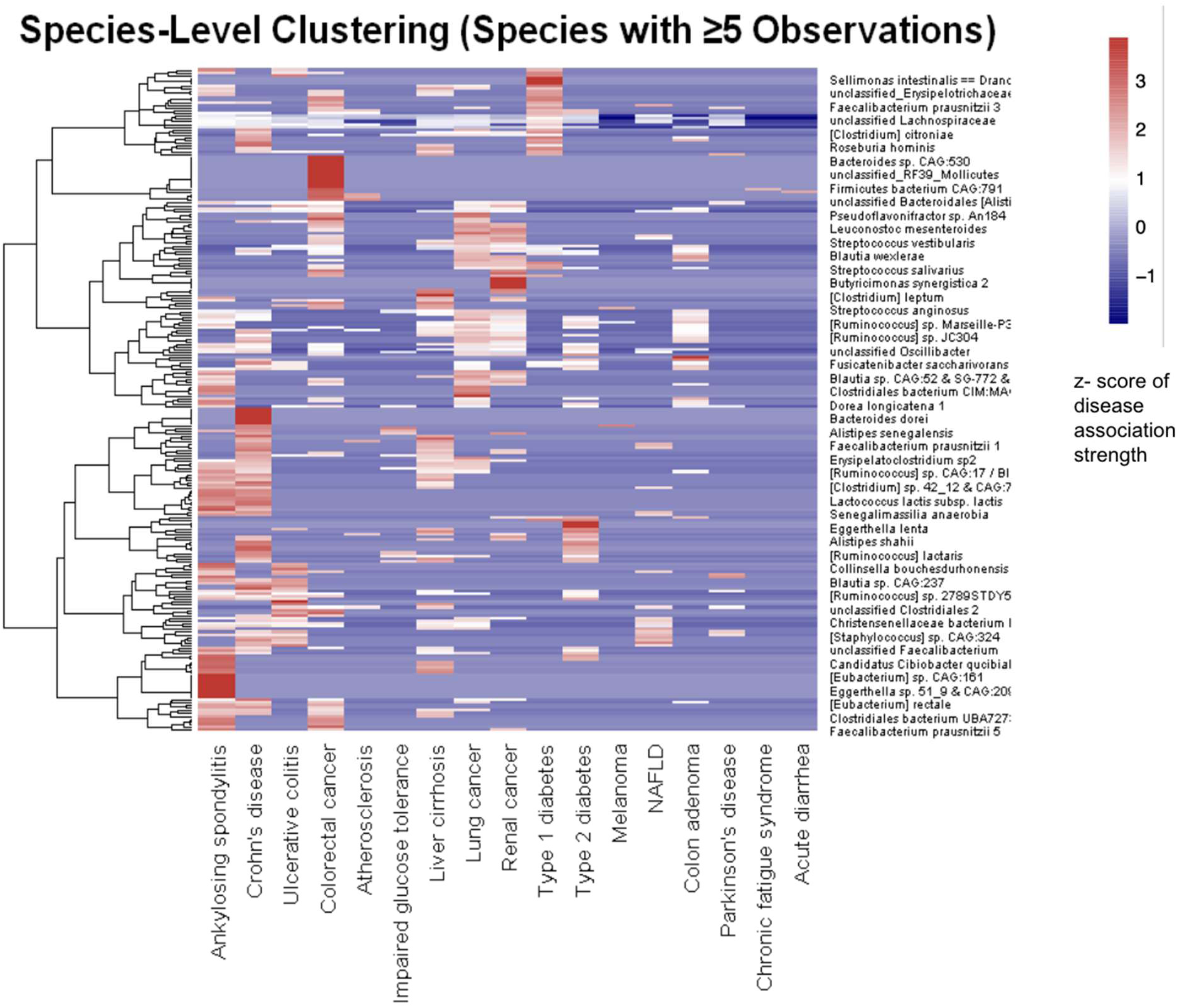
Hierarchical clustering heatmap. *Figure 1 illustrates the relationship between Microbial Species (count greater than or equal to 5) and their Disease Association. Color intensity represents the score of disease association strength with dendrograms indicating similarity in microbial signatures across different diseases and species clusters.*

**Table 4:** Relationship between disease and bacteria based on red classification.

| *Disease* | *Bacteria* |
| --- | --- |
| Ankylosing Spondylitis | [Eubacterium] sp. CAG: 161 |
| Ankylosing Spondylitis | Eggerthella sp 51_9 & CAG:209 |
| Crohn's Disease | Bacteroides dorei |
| Ulcerative Colitis | [Ruminococcus] sp. 2789STDY5 |
| Colorectal Cancer | Bacteroides sp. CAG:530 |
| Colorectal Cancer | unclassified_RF39_Mollicutes |
| Liver Cirrhosis | [Clostridium] leptum |
| Lung Cancer | Clostridiales bacterium CIM:MA |
| Renal Cancer | Butyricimonas synergistica 2 |
| Type 1 Diabetes | unclassified_Erysipelotrichacea |
| Type 2 Diabetes | Eggerthella lenta |
| Colon adenoma | Unclassified Oscibillibacter |

**Table 5:** Distribution amongst countries.

| ***Cluster Label*** | ***Data Points*** | ***Top Countries*** | ***Top 2 Diseases*** | ***Top 2 Genera*** | ***Disease Association Range*** |
| --- | --- | --- | --- | --- | --- |
| ● **Noise** | 126 | AUT, FRA, TZA | Colorectal cancer, Renal cancer | Prevotella, Lachnoclostridium | 0.308-0.651 |
| ● **0** | 2197 | SWE, FRA, PER | Colorectal cancer, Liver cirrhosis | Blautia, unclassified Lachnospiraceae | 0.300-0.538 |
| ● 1 | 13 | ESP, AUT, CHN | Melanoma, Liver Cirrhosis | Lachnoclostridium | 0.307-0.359 |
| ● 2 | 5 | AUT, DEU | Lung cancer, Renal cancer | Blautia, Coprococcus | 0.555-0.594 |
| ● 3 | 109 | PER, TZA, MNG | Colorectal cancer, Liver cirrhosis | unclassified Clostridiales 3, unclassified Clostridiales 5 | 0.300-0.468 |
| ● 4 | 12 | DNK, GBR, FRA | Liver cirrhosis, Colorectal cancer | unclassified Clostridiales Family XIII. Incertae Sedis, unclassified Clostridiales 5 | 0.381-0.433 |
| ● 5 | 11 | FRA, GBR, ESP | Lung cancer, Renal cancer | Blautia, Anaerostipes | 0.572-0.605 |
| ● 6 | 8 | DEU, ESP, AUT | Lung cancer, Colorectal cancer | Flavonifractor, Lachnoclostridium | 0.495-0.529 |
| ● 7 | 4 | ITA, JPN LUX | Lung cancer, Colon adenoma | Flavonifractor, Lachnoclostridium | 0.526-0.529 |
| ● 8 | 5 | SWE, MNG, THA | Liver cirrhosis, Colon adenoma | Roseburia, Ruminiclostridium | 0.318-0.349 |
| ● 9 | 14 | SWE, THA, MNG | Lung cancer, Renal cancer | Coprococcus, Dorea | 0.533-0.594 |

### 4.2 DATA VISUALIZATIONS

Figure 2 visualizes the distribution of microbial species counts across countries and disease categories using datasets from the Human Gut Microbiome Atlas. The bars represent cumulative species counts per country, with individual segments corresponding to specific disease conditions. The visualization highlights notable geographic variability in microbial diversity, with countries such as Sweden, France, and Denmark exhibiting higher overall species counts compared to countries including the United States, India, and China. Differences in disease-specific contributions to total species counts are also apparent, suggesting heterogeneous representation of disease-associated microbial taxa across geographic regions.

**Figure 2:**
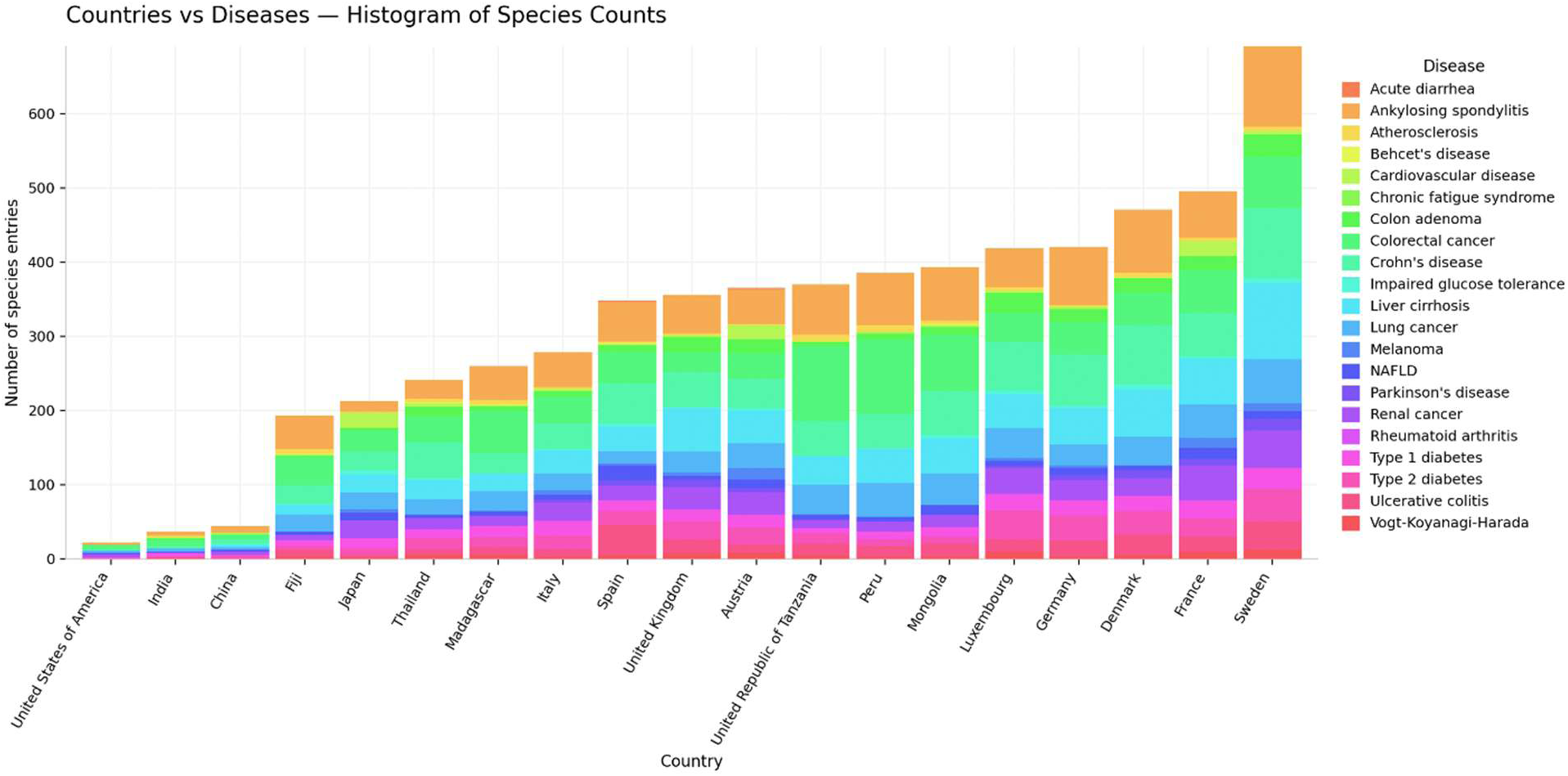
Histogram of Species Counts per country. Figure 2 represents the total number of unique microbial species entries identified per country. The x-axis identifies 19 distinct countries, while the y-axis quantifies the number of species entries. The colors in the histogram indicate the proportional contribution of specific diseases (e.g., Crohn’s disease, Type 2 diabetes, Liver cirrhosis) to the microbial origin of each country.

Figure 3 presents mean effect size (ES) values for multiple disease conditions across countries based on Human Gut Microbiome Atlas data. Each bar reflects the aggregated effect score per country, summarizing the magnitude of disease-associated microbial shifts across conditions. Higher mean effect sizes are observed in countries such as Austria, Spain, and France, whereas lower values are seen in countries including the United States and India. This visualization displays an international comparison of disease-related microbiota.

**Figure 3:**
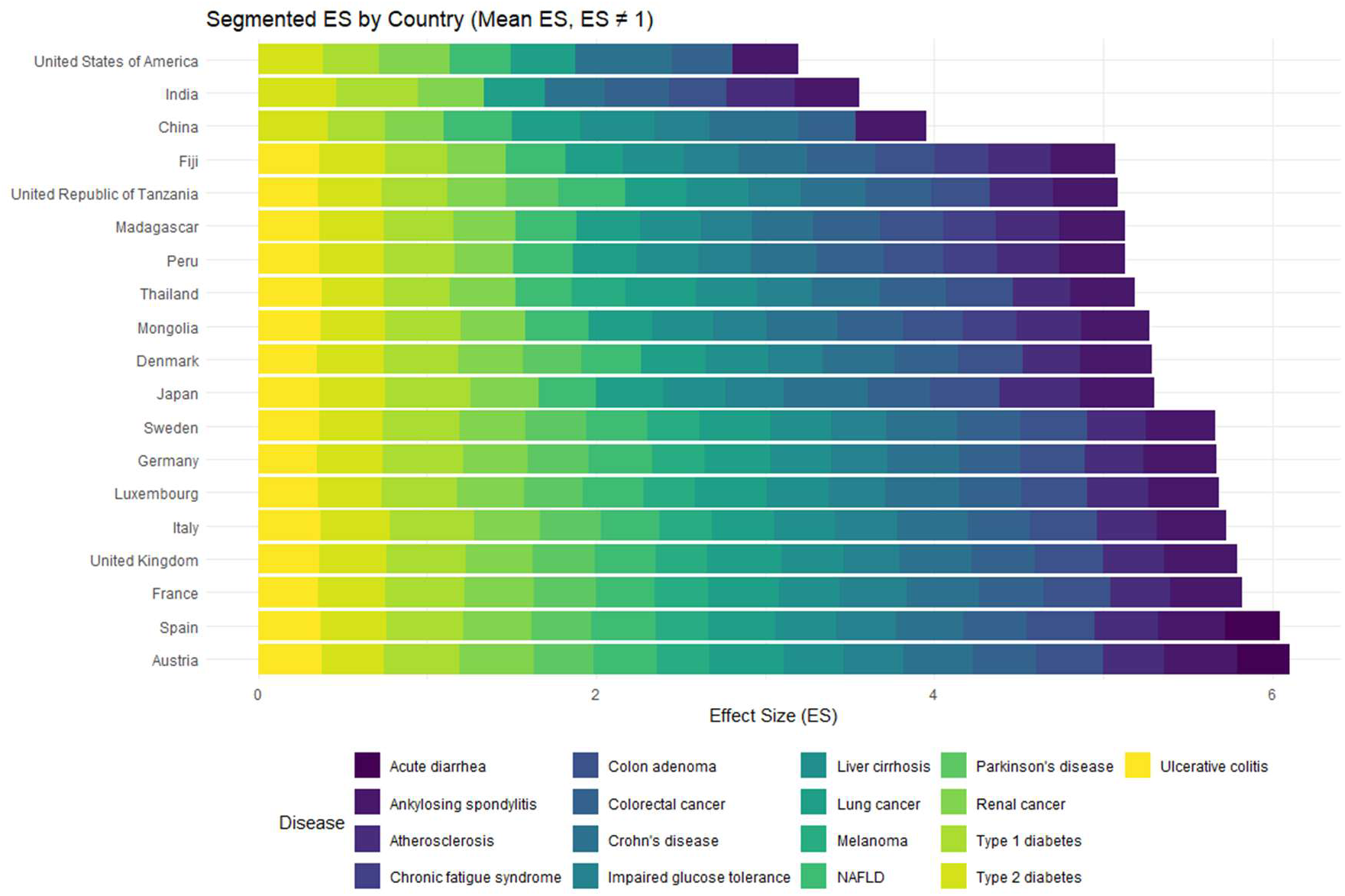
Bar Chart Depicting Country ES (Effect Score) *Figure 3 quantifies the strength of association (Mean ES) between the gut microbiome and 19 distinct countries. The multi-colored bars illustrate the distribution of effect sizes across different diseases, highlighting countries with the highest aggregate microbial values*.

### 4.4 MACHINE LEARNING

The comparison of machine learning model performances across microbiome datasets reveals that ensemble-based tree models, particularly Bagging, Boosting, and Random Forest, consistently achieved the highest classification accuracies across populations and disease types. Westernized datasets generally produced higher accuracies than non-Westernized datasets, reflecting differences in the similarity of microbiomes within each population. Overall, these results highlight the effectiveness of ensemble-based tree methods in capturing patterns within microbiome data.

#### 4.4.1 DBSCAN CLUSTERING

Figures 4 illustrates microbial patterns within the cancerous dataset, where there is a significantly higher taxonomic diversity compared to westernized and non-westernized cohorts. The DBSCAN plot identifies 10 distinct clusters nested within the dense dataset. Cluster 0 serves as a broad, baseline population with 2197 data points, and other smaller clusters (Clusters 2, 5, and 9) are grouped with high Disease Association Scores and contain specific genera such as Coprococcus and Biautia.

**Figure 4:**
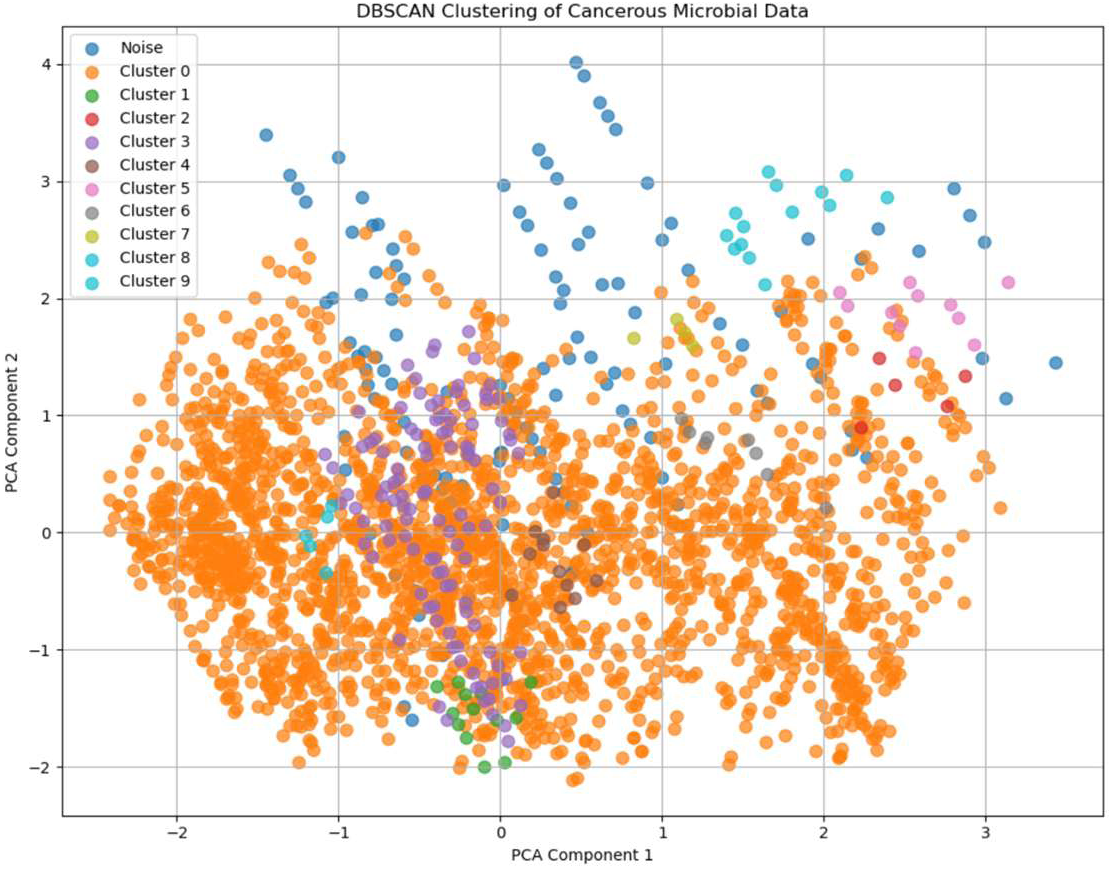
Principal component based on disease. *Figure 4 demonstrates the DBSCAN (Density-Based Spatial Clustering of Applications with Noise) applied to the first two principal components of the cancerous microbial dataset. Points are color-coded by identified clusters (0–9) with blue points representing statistical noise*.

#### 4.4.2 KNN

As Table 6 shows, KNN has relatively low and inconsistent accuracy across the datasets, making it less reliable than other models. Additionally, the KNN machine learning model was applied to all datasets and showed clear differences among them. The westernized dataset achieved 24% accuracy, while the non-westernized dataset reached 31%, suggesting that the non-westernized data is more consistent and easier to distinguish for disease classification. The cancerous dataset produced the highest accuracy at 41%. Overall, the cancerous dataset behaved differently from both the westernized and non-westernized datasets across the machine learning models. These low accuracies showed that KNN was not a viable ML technique for our dataset so we discontinued it when applying ML techniques to the cancerous westernized and cancerous non-westernized datasets.

**Table 6:** Accuracy for KNN.

| *Model* | *Accuracy: Cancerous* | *Accuracy: Westernized* | *Accuracy: Non-westernized* |
| --- | --- | --- | --- |
| KNN | 41.0% | 24.0% | 31.0% |

#### 4.4.3 RANDOM FOREST

From Table 7, it is clear that Random Forest achieved its highest accuracy in predicting cancerous cases in Westernized populations, reaching 91.0% for cancerous Westernized samples and 76.3% for cancerous non-Westernized samples. For non-cancerous cases, the model reached 87.0% accuracy in Westernized populations and only 72.9% in non-Westernized populations. Across all categories, the highest accuracy was observed for cancerous Westernized samples, while non-cancerous non-Westernized samples were the lowest, indicating a consistent performance gap of approximately 15-16% between Westernized and non-Westernized populations.

**Table 7:** Accuracy across ML Models.

| *Model* | *Accuracy: Cancerous Westernized* | *Accuracy: Cancerous Non-Westernized* | *Accuracy: Westernized* | *Accuracy: Non-westernized* |
| --- | --- | --- | --- | --- |
| Random Forest | 91.0% | 76.3% | 87.0% | 72.0% |
| Bagging | 91.0% | 84.0% | 92.0% | 78.0% |
| Gradient Boosting | 91.5% | 84.4% | 89.4% | 72.2% |
| Neural Networks | 50.8% | 52.6% | 32.1% | 36.0% |

#### 4.4.4 BAGGING AND GRADIENT BOOST

As Table 7 demonstrates, Bagging achieved the second-highest accuracy across all datasets. However, Boosting consistently outperformed it, and when Boosting was performed on the different types of group its accuracy was always the highest. Therefore, boosting methods, particularly Gradient Boosting, produced the strongest consistent results, achieving peak accuracies of 91.5% for the cancerous westernized dataset. Although it did have a low of 77.2% for the non-westernized dataset, that was the highest accuracy for the non-westernized dataset across all ML methods.

Neural Network performance is consistently weak, as Table 7 demonstrates. These results indicate poor classification capabilities of the algorithm. Specifically, the model achieved an accuracy of only 50.8% for cancerous Westernized samples and 52.6% for cancerous non-Westernized samples. For non-cancerous cases, the accuracy was as low as 32.1% in Westernized populations and 36.0% in non-Westernized populations. Overall, the model performed substantially lower than the other models across every category. Furthermore, the gap between the accuracies across each ML algorithm is significant, making it the least effective model in this comparison.

### 4.5 DIMENSIONALITY REDUCTION

Dimensionality reduction revealed the difference in relation to each disease. As shown in Figures 5a and 5b, the UMAP and tSNE projections displayed a lack of a distinct isolated group, and instead, the dotted variance of color across the plots indicates that different diseases share many overlapping features unable to be distinguished by these identifiers alone. Rather than unique microbe-to-disease relationships, different diseases are driven by a shared state of dysbiosis. The data appear in stringy shapes, suggesting a gradient between different disease states.

**Figure 5a:**
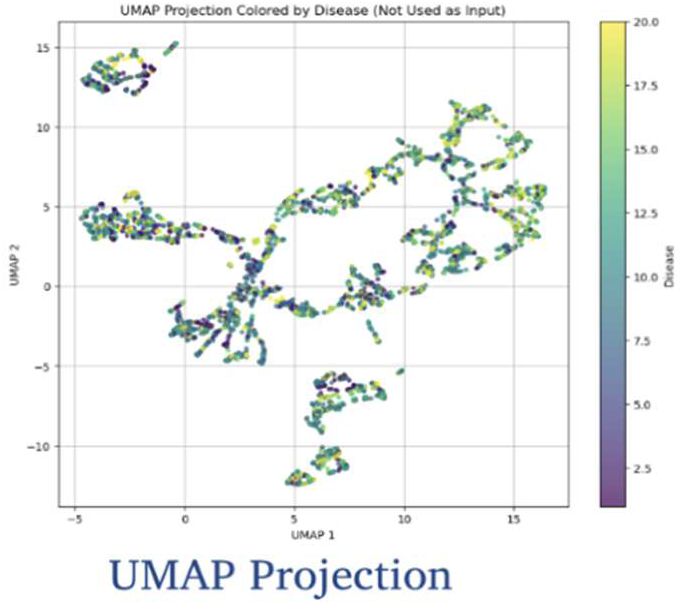
Dimensionality reduction UMAP. *Figure 5a displays a projection of the dataset after employing UMAP for dimensionality reduction. The plot points are colored by disease to reveal how diseases have similar or different effects on human gut microbiome composition*.

**Figure 5b:**
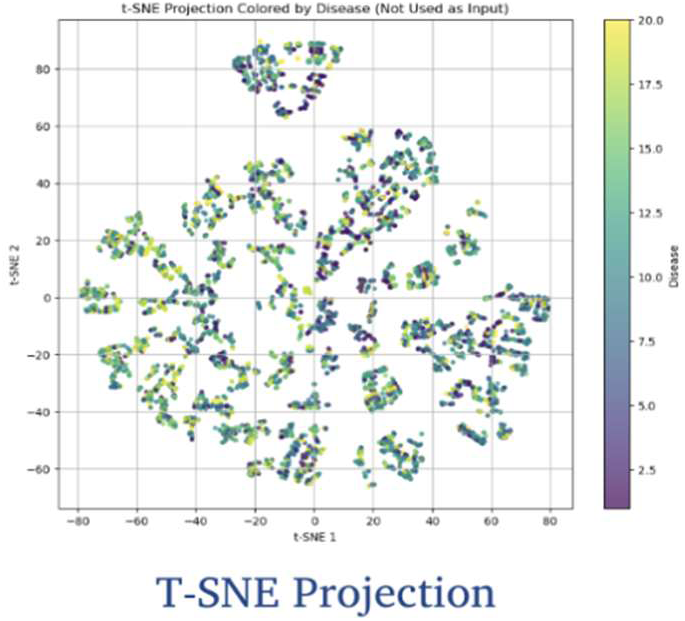
Dimensionality reduction t-SNE. *Figure 5b displays a projection of the dataset after employing T-SNE for dimensionality reduction. The plot points are colored by disease to reveal how diseases have similar or different effects on human gut microbiome composition*.

### 4.6 TOPOGRAPHICAL DATA ANALYSIS

Figure 6 shows the Mapper visualization’s changes when the chosen resolution parameter *n_cubes* is changed. The overlap parameter was kept fixed at 0.3, and DBSCAN clustering was used for all to ensure any changes were attributed solely to the number of intervals. At lower resolutions (*n_cubes* = 5 and 10), the resulting graphs are compact and densely connected, capturing the central global structure of the dataset. In these cases, DBSCAN produces relatively large clusters that span multiple regions, leading to simplified topological representations. These configurations, despite preserving connectivity, also provide a more limited insight into variations in underlying patterns within the data.

**Figure 6:**
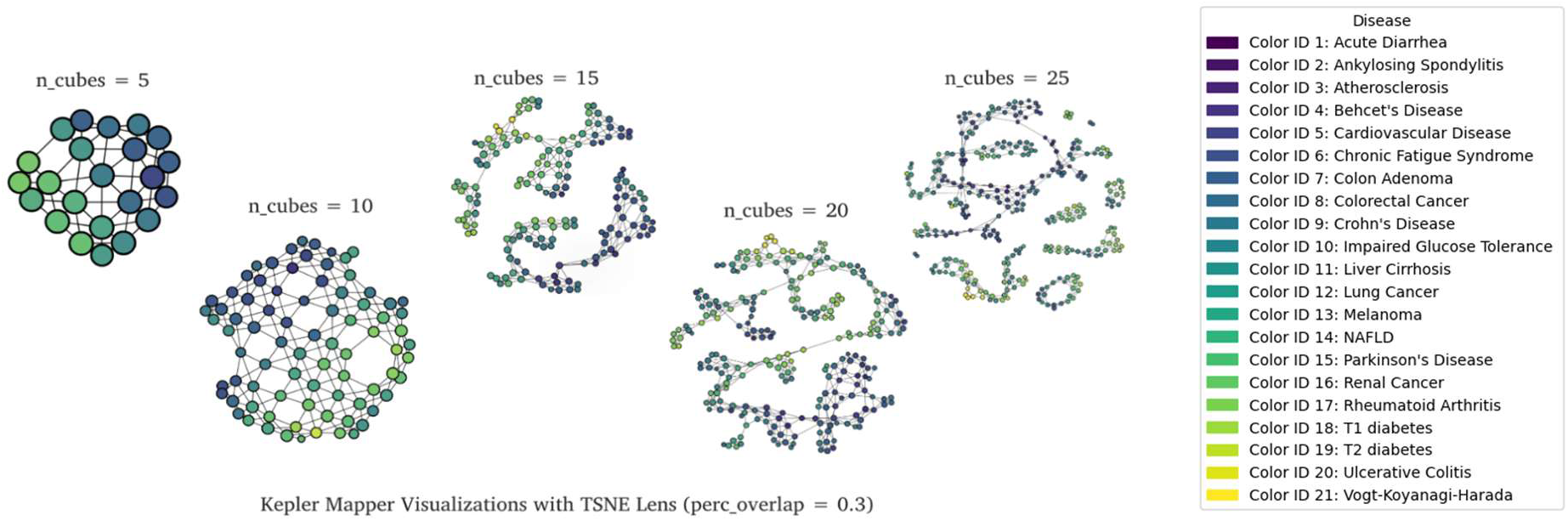
Topographical Data Analysis. *Figure 6 reveals the global and hidden structures within the Human Gut Microbiome dataset through a series of Kepler Mapper visualizations. This figure highlights how increasing resolution parameters, specifically the number of n_cubes, impacts the clustering of data points by creating more complex structures*.

This hidden structure is more clearly revealed through the Kepler Mapper visualizations. As the lens resolution increases from *n_cubes = 5* to *n_cubes = 25*, the visualization shifts from one solid mass into a detailed network of loops and branches. These branches represent points of divergence where certain subsets of data begin to behave differently from the main group, while the loops display repeating patterns that occur across different categories.

Increasing the resolution (*n_cubes* = 15 to 25) results in more complex Mapper graphs that have greater fragmentation and branching. At these higher values, smaller and more localized clusters are identified, which produces elongated and partially disconnected structures. This suggests that although higher resolutions perform better in revealing subtle subclusters, these benefits come at the expense of overcomplication of actual patterns in the data. These findings indicate a trade-off between interpretability and detail in the structures, with higher values of *n_cubes* offering increased sensitivity to understated patterns at the expense of possibly wrongly identifying non-existent clustering, while not having enough intervals can result in oversimplification of the data’s structure.

## 5. DISCUSSION

### 5.1 GLOBAL DISEASE AND MICROBIOME ASSOCIATION PATTERNS

Disease association scores across countries vary based on the categorical values of country, species, genus, and disease, as demonstrated by ANOVA visualizations. The ANOVA results indicate that biological variables explain substantially more variation in Disease Association than geographic origin. Species level explains the greatest total variance within the dataset, demonstrating that disease association patterns differ strongly at the species level. Following this, the Disease category shows the highest F value, demonstrating big differences in association scores across diseases. Additionally, Genus also significantly influences Disease Association to a lesser extent to species level. In contrast, Country shows no significance in the variation of Disease Association, indicating that geographic origin does not structure association scores in this dataset. Although several predictors are highly statistically significant, a large portion of the variance remains unexplained.

### 5.2 SHARED DYSBIOSIS PATTERNS ACROSS DISEASES

UMAP and t-SNE show overlapping disease samples, with no clear disease-specific clusters. Color gradients demonstrate diseases sharing dysbiosis patterns rather than distinct signatures. UMAP shows gradual global transitions while t-SNE highlights local clusters with mixed diseases. These patterns show that diseases share similar disruptions in the microbiome rather than having distinct configurations.

### 5.3 KEPLER MAPPER ANALYSIS

Kepler Mapper displays hidden structures within the gut microbiome data as the gradients show mixed disease labels in nodes, indicating shared disease dysbiosis. Low resolution (*n_cubes* = 5) oversimplifies the topology, while high resolutions (*n_cubes* = 15 – 25) fragments the graph due to noise. Thus, *n_cubes* = 10 best balances the factors, demonstrating the shared dysbiosis present in the dataset. Alongside the branches and loops that display gradual transitions between microbial states, the diseases overlap instead of forming distinct clusters, showing clear variation in the dataset. Using the legend from the TDA visualization, disease patterns were identified within the network. There is a cluster of inflammatory diseases, including Crohn’s disease, ulcerative colitis, and rheumatoid arthritis, which overlap with one another. Cancer-related diseases, such as colorectal cancer, colon adenoma, lung cancer, and melanoma, are also spread across multiple clusters, but these clusters are more fragmented. This indicates a greater level of complexity in cancer-related microbiomes. Metabolic disorders such as diabetes type I and II and impaired glucose tolerance show partial overlap with one another. Liver disease, such as NAFLD and liver cirrhosis, is also clustered in a similar area of the network. However, neurological and systemic diseases such as Parkinson’s, multiple sclerosis, and chronic fatigue syndrome are more distinct from one another. This indicates that there is a greater level of variation in the microbiome for these types of disease. While there is a level of overlap in the microbiome for certain types of disease, there is also a level of distinctness.

### 5.4 CANCEROUS VS NON-CANCEROUS MICROBIOME PROFILES

The DBSCAN results show greater heterogeneity in cancerous microbiomes, which have more clusters and a wider range of disease association values in comparison to non-cancerous microbiomes. In the cancerous dataset, Clusters 1-9 have disease association values from approximately 0.30 to 0.60, reflecting variation in microbial composition. For example, lower-range clusters such as Cluster 1 (0.30-0.36) are dominated by Clostridium species, while higher-range clusters such as Cluster 5 (0.57-0.60) and Cluster 2 (0.55-0.59) include genera like Blautia, Coprococcus, and Anaerostipes.

In contrast, the non-cancerous data contains tighter clustering with fewer outliers. The DBSCAN plot for non-cancerous diseases has a single large cluster indicated by Cluster 0, n=323, and therefore is not insightful nor relevant in this study.

### 5.5 PERFORMANCE OF MACHINE LEARNING MODELS

For all categories of datasets, ensemble-based tree data models showed the highest classification accuracy. Boosting-based techniques were found to possess the strongest and most consistent performances among all techniques. Peak accuracy rates were achieved by these techniques in westernized and cancerous datasets, with rates over 90%. These techniques were able to manage the high dimensionality of the data well.

### 5.6 SHARED DYSBIOSIS AND DISEASE PREDICTION IMPLICATIONS

Overlapping microbiome signatures across diseases imply that prediction models may be predicting common dysbiosis states rather than disease-specific microbial profiles. Population-specific performance of the prediction models emphasizes the need to consider geographic and lifestyle factors.

### 5.7 CLINICAL AND RESEARCH SIGNIFICANCE

Identifying shared dysbiosis patterns across diseases may improve understanding of common pathological mechanisms, particularly in gut–brain axis disorders. The integration of geographic information enhances the interpretability and generalization of microbiome-based disease models.

## 6. LIMITATIONS

### 6.1 STUDY LIMITATION

Despite the strong performances of the ensemble-based tree models, differences in sample size across groups and relatively small cohort sizes may have affected the stability and accuracy of each model, particularly for diseases with fewer samples. For instance, the KNN model showed low accuracy across datasets due to its reliance on comparing samples directly to each other, which would not work well when the data is complex and has overlapping microbial patterns across diseases.

### 6.2 LIMITATIONS OF DISTANCE-BASED AND NEURAL NETWORK MODELS

The KNN model displayed low classification accuracies across categories, indicating limitations in distance-based ML methods for complex microbiome data. Confusion matrices showed significant misclassification, specifically, the reduced performance in westernized datasets, suggesting that overlap in microbial features across diseases and populations makes it harder for models to tell them apart. Neural Networks demonstrated the weakest performance, with accuracy close to random predictions, likely due to class imbalance, limited sample numbers compared to the large number of microbial features, and the overall complexity of the data. Similarly, NN showed a tendency to generate high confidence predictions for underrepresented categories, which further increased misclassification.

## 7. FUTURE WORKS

Future work incorporating functional genomics, expanded cohorts, and external validation may further enhance the clinical relevance and generalizability of microbiome-driven predictive models. Expanding datasets to include functional and multi-omics information could improve model interpretability and biological relevance. Increasing sample sizes, especially for underrepresented diseases, may improve neural network and distance-based model performance. Additional validation using independent cohorts is necessary for assessing model robustness and ensuring reproducibility across populations. Further exploration of topological methods may uncover higher-order structures in microbiome data that are relevant to disease progression and population-level variation. These insights highlight the potential of gut microbiome research to inform disease prevention, early detection, and personalized healthcare strategies. Continued studies building on these findings could transform our understanding of microbiome-disease interactions on a global scale. Expanding on this work could enhance our ability to connect gut microbiome profiles with health outcomes, enabling more precise interventions and preventative care.

## 8. CONCLUSION

This study aimed to apply complex data analysis and machine learning techniques to improve the accuracy of prediction algorithms for identifying gut-related diseases and microbiome associations. By integrating ensemble-based machine learning methods with TDA, our work demonstrates that relationships between microbiota and diseases are driven by nonlinear interactions rather than simple taxonomic or geographic labels. At a higher resolution (n_cubes = 25), several nodes show clear dominance of specific disease-associated colors from the legend, particularly those representing type 2 diabetes, Crohn’s disease, and colorectal cancer. These patterns indicate localized enrichment of these conditions, although the nodes remain partially connected to neighboring regions containing mixed disease categories. This suggests that while certain diseases display more concentrated microbiome signatures, they are still embedded within broader, shared dysbiosis patterns. By analyzing the complex, high-dimensional, and sparse nature of gut microbiome data, the methods can also improve the classification and prediction accuracy of various diseases.

The results demonstrate that tree-based ensemble models, specifically Bagging and Boosting, outperform other machine learning techniques in identifying patterns and clusters in the data used in this project due to their ability to better handle more complex, multi-dimensional data without overfitting, with Gradient Boosting reaching up to 91.5% accuracy. These findings highlight the strong potential of computational approaches such as tree-based ensemble models to reveal meaningful patterns in the gut microbiome that were previously difficult to detect. The strongest disease associations among cancers, specifically lung and renal cancers, are taxonomically defined by enrichment for the genera Blautia, Coprococcus, and Dorea. In contrast, the study identified pathogenic markers of Fusobacterium nucleatum and Flavonifractor plautii as they were isolated in their own environments. The research underscores the broad applicability of microbiome insights across multiple diseases, emphasizing the value of understanding shared dysbiosis patterns for future clinical and therapeutic strategies.

In addition, the study’s conclusions indicate geographic stratification, with European countries such as Sweden and Austria comprising the majority of the dense baseline and high-cancer association clusters, consistent with other studies that attribute this to characteristics of advanced healthcare systems and long life expectancy. Samples from Peru, Tanzania, and Mongolia are grouped together and show strong evidence that biogeographic factors and disease states influence their microbial structure. This can help guide future studies toward identifying biomarkers and improving disease prediction, ultimately supporting personalized medicine approaches.

## AI ASSISTANCE DISCLOSURE

An AI-powered writing assistant was used solely to refine grammar and spelling. No generative artificial intelligence was used to create data values or generate text for the manuscript.

## ACKNOWLEDGEMENTS

This research received no specific grant from any funding agency in the public, commercial, or not-for-profit sectors.

The authors would like to thank the Aspiring Scholars Directed Research Program and the Olive Children Foundation for providing this opportunity and making this study possible. In addition, the authors would also like to extend their gratitude to the Principal Investigator, Sahar Jahanikia, for her invaluable support and guidance.

## DATA, METADATA, AND CODE AVAILABILITY STATEMENT

The metadata supporting the results obtained through this is based on the Human Gut Microbiome Atlas dataset, which is publicly accessible. These consist of microbial taxonomy data (species level, genus level), geographical information (country), diseases, and disease association scores. The metadata for the study will be deposited in an appropriate repository platform after peer review.

The documentation of the metadata contains descriptions of the preprocessing methods, definitions of the variables, and the format of the datasets. In this study, the authors adhered to the guidelines recommended by the STORMS (Strengthening the Organization and Reporting of Microbiome Studies).

All the code used for the data pre-processing, statistical analyses, machine learning model training, and visualization performed in this research will be openly shared through a version-controlled repository and archived using a DOI once the paper is published.

## DATA AVAILABILITY STATEMENT

All data used in this study are publicly available through the Human Gut Microbiome Atlas (HGMA), which provides global metagenomic data including species and genus-level abundance, disease associations, and geographic distributions across 20 countries. The HGMA database is openly accessible at https://www.microbiomeatlas.org.

